# TrEMOLO: Accurate transposable element allele frequency estimation using long-read sequencing data combining assembly and mapping-based approaches

**DOI:** 10.1101/2022.07.21.500944

**Authors:** Mourdas Mohamed, François Sabot, Marion Varoqui, Bruno Mugat, Katell Audouin, Alain Pélisson, Anna-Sophie Fiston-Lavier, Séverine Chambeyron

**Affiliations:** Institute of Human Genetics, UMR9002, CNRS and Université de Montpellier, Montpellier, France; DIADE, University of Montpellier, CIRAD, IRD, Montpellier, France; IFB - Southgreen Bioversity, CIRAD, INRAE, IRD, Montpellier, France; TAGC, UMR 1090 INSERM, Marseille, France; ISEM, Université Montpellier, CNRS, IRD, CIRAD, EPHE, Montpellier, France; Institut Universitaire de France (IUF)

**Keywords:** Genome, software, transposable element calling, long-read DNA sequencing, nanopore sequencing, transposable element allelic frequency, haplotypes, structural variation

## Abstract

Transposable Element MOnitoring with LOng-reads (TrEMOLO) is a new software that combines assembly- and mapping-based approaches to robustly detect genetic elements called transposable elements (TEs). Using high- or low-quality genome assemblies, TrEMOLO can detect most TE insertions and deletions and estimate their allele frequency in populations. Benchmarking with simulated data revealed that TrEMOLO outperforms other state-of-the-art computational tools. TE detection and frequency estimation by TrEMOLO were validated using simulated and experimental datasets. Therefore, TrEMOLO is a comprehensive and suitable tool to accurately study TE dynamics. TrEMOLO is available under GNU GPL3.0 at https://github.com/DrosophilaGenomeEvolution/TrEMOLO.

## Background

Transposable Elements (TEs) are endogenous mobile elements that can move within their host genome and increase their copy number, autonomously or not (1). They are present in almost all genomes that have been analyzed, and are represented by various structures and mobility mechanisms. Their size ranges from less than 100 bp (e.g. for MITEs (2)) to more than 20kb (3,4). In large genomes, they represent more than 90% of nuclear DNA (5), but they whatever are a major component of smaller genomes. For instance, the *Drosophila melanogaster* genome is only 180Mb long, but almost ~15% of it is made of TEs (6). However, their identification, annotation, and dynamic tracking are complex due to their intrinsic repetitive nature. Indeed, identifying insertions in a newly assembled genome is a hard and complex task. Many tools have been developed to tackle this issue (*e.g*., REPET (7), RepeatMasker (8), EDTA (9)), with more or less success, and requiring all a large amount of computational resources. Moreover, tracking the presence or absence of a given insertion among individuals or through generations was, up to recently, limited to heavy wet-laboratory approaches, with a low throughput (10–12).

Sequencing technology advances in the last 15 years, especially Illumina short-read sequencing, have allowed sequencing, on a daily basis, not only individuals within a species or subspecies, but also within whole populations. Nowadays, it is not uncommon to sequence the same population many times over generations to study its genetic evolution (13). One of the main findings provided by the availability of massive sequencing data is that TEs are the most variable component of genomes, even more than previously suspected (14). Therefore, many tools have been developed to use short-read sequencing data to identify potential TE polymorphisms among samples (*e.g*., T-lex (15–17); McClintock (18)) or within a population (*e.g*., DNApipeTE (19); PoPoolationTE2 (20)). They rely on three main mapping features: split reads, unpaired reads, and depth of coverage (14). However, short-read sequencing data are not suitable for the optimal detection of structural variations (SV), especially SV due to TE insertion/deletion. Indeed, as short-reads are shorter (100 to 250 bases) than the mean length of repeats and TEs, they do not span the entire length of the variation and thus cannot be used to correctly identify SV, leading to high false-positive rates (40%) and false-negative results (21,22). Conversely, Long Read (LR) sequencing technologies, such as Pacific Biosciences and Oxford Nanopore Technologies (ONT), allow obtaining reads that are >20kb long (and even 100kb and more for ONT). Therefore, a single read may contain a complete SV or TE insertion and enough surrounding sequence data to unambiguously anchor the variation. However, the longer read length is counterbalanced by a lower sequencing quality (still improving) compared with short-read sequencing and fewer dedicated tools for TE detection. The current methods to detect TE variations using LR sequencing data (and also most of the short-read sequencing data-based tools) generally rely on the presence/absence of TE insertions annotated in a reference genome (*e.g*., LorTE (23)), or on the detection of abnormally mapped reads upon the reference genome that can be linked to a TE (*e.g*., TELR (24)). However, some tools also allow the partial detection, assembly and annotation of non-reference TE insertions (e.g. TLDR) (Transposons from Long DNA Reads) (25).

In addition, unlike short-read, LR sequencing allows generating high-quality genome assemblies with higher levels of contiguity, and consequently to compare genomes directly and to extract more efficiently large and complex SVs (26). Indeed, the number of available almost complete genomes is increasing due to these LR sequencing technologies (13,27), and even telomere-to-telomere assemblies can be obtained (28). Moreover, some reliable tools are available for genome-to-genome comparisons (MUMmer (29), minimap2 (30)) and for identifying complex SV between two assemblies more efficiently than with mapping-based approaches. However, assembly methods generally rely on a layout-and-consensus step when solving the multiple possibilities at a given locus (*i.e*., choosing one of all possible haplotypes in the graph, usually the most represented one), and thus may mask some minor populational/somatic variations.

In this paper, we present the Transposable Element MOvement detection using LOng-reads (TrEMOLO) software that combines the advantages offered by LR sequencing (*i.e*., highly contiguous assembly and unambiguous mapping) to identify TE insertion (and deletion) whatever the frequency at which they segregate in a population. TrEMOLO (available at https://github.com/DrosophilaGenomeEvolution/TrEMOLO) is a SnakeMake pipeline that can be used in script mode with dependencies already installed, or within a Singularity container to ensure a higher reproducibility and to facilitate future updates. The pipeline is designed so that the inclusion of additional samples is straightforward. We demonstrated its efficiency using simulated and experimental LR sequencing data from a highly polymorphic *Drosophila* population. TrEMOLO detected TE insertions that could not be identified with other approaches, with a very low error rate. Therefore, TrEMOLO provides an unprecedented accurate picture of TE variability compared with other tools.

## Results

### TrEMOLO design

TrEMOLO is a suite of Python 3 (31) scripts that follow the SnakeMake (32) rules. It takes as input a TE library, a reference genome sequence, single or pooled LR DNA sequencing data, and a genome assembly generated from these sequencing data. In the sequel, TE insertions from the major haplotype belonging to the assembly will be called INSIDER TE insertions. The minor frequency haplotype TE insertions are not included in the assembled genome and will be called OUTSIDER TE insertions. The complete tool includes three internal modules (Fig.1 and Fig. S1): (1) the ***INSIDER TE detection module*** identifies TE insertions and deletions present only in the newly assembled genome when compared to the reference genome, by aligning these two genomes and parsing the pairwise alignment; (2) the ***OUTSIDER TE detection module*** searches for the additional TE insertions (those that are carried by rare haplotypes and are therefore not present in the assembly), by mapping, LRs to the genome assembly generated using the same data, and detecting TE variants by parsing the mapping results (Fig.1 and Fig.S1); (3) the ***TE analysis module*** computes from the previous modules outputs the frequency of each TE variant using the LR sequencing data and looks for target site duplications (TSDs) that occur upon TE integration as a result of staggered DNA double-strand breaks (33). A detailed manual on how to run TrEMOLO with all its options is available at TrEMOLO github (https://github.com/DrosophilaGenomeEvolution/TrEMOLO).

**Figure 1.**
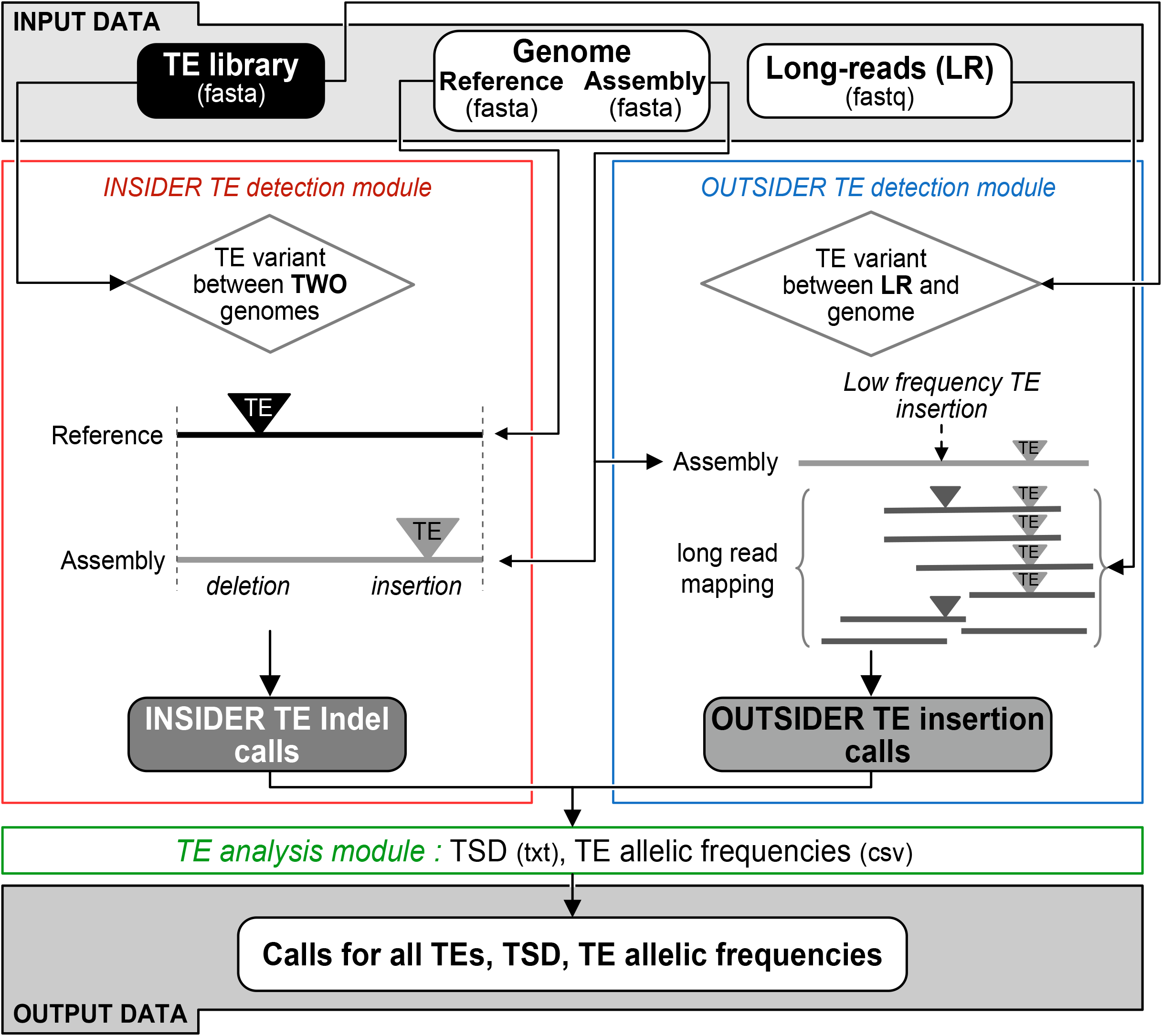
Overview of the TrEMOLO pipeline. Three main modules: (1) the INSIDER TE detection module in red, (2) the OUTSIDER TE detection module in blue, and (3) the TE analysis module in green.

#### The INSIDER TE detection module

This module detects the so-called INSIDER TEs. These are the non-reference TE insertions/deletions that are found frequently enough among the reads to be incorporated in the assembly (generally the TE polymorphisms of the major haplotype). It takes as input a TE library, a reference genome, and the genome assembly of the studied biological sample, all in fasta format. This assembly must be generated with the input LR sequencing data to be compatible with the second module. Whole-genome pairwise alignment is performed between the reference and assembled genomes using minimap2 (30), and is then parsed with *Assemblytics* (34) for SV identification (Fig. S1). This analysis returns a list of SVs from the pairwise alignment of the assembly against the reference genome with their positions. This list is then compared with a TE library to keep only SV belonging to already known TE families (Fig. 1, left panel, and Fig. S1). Before providing the final list of detected TE insertions/deletions, the alignment data are scanned to filter out sequences that do not fit the parameters (considered as low-quality TEs), corresponding here to 80% of the size and 80% of identity (Methods).

#### The OUTSIDER TE detection module

This module allows to retrieve OUTSIDER TEs, i.e. those TE polymorphisms that are low frequency in the sample and not included in the genome assembly. This module takes as input a TE library in fasta format, a genome assembly (fasta) and the LR sequencing data in fastq used to build this assembly. The reads are aligned to the assembled genome using minimap2 (30). Reads that are flagged as partially or non-mapping in the output SAM file are candidates for carrying OUTSIDER TEs. Candidate reads are validated if the corresponding sequences are present in the provided TE library (Fig. 1 right panel, Fig. S1). As for the INSIDER module, low-quality TEs are filtered out and then, TE insertions are listed in a BED file.

#### The TE analysis module

This module is dedicated to the analysis of the newly inserted TEs (a) to detect hallmarks of the transposition mechanism (*i.e*. TSDs) and (b) to estimate the allele frequency of each TE insertion.

(a) For the INSIDER TE insertions, the 30 bp flanking both extremities of the TE insertion are aligned against each other using the Boyer-Moore exact algorithm (35) to detect short motifs shared on each side of the insertion without mismatches. For the OUTSIDER TE insertions, the most informative single LR (with the higher BLAST score) spanning the TE insertion is selected. Then, like for INSIDER TE insertions, the sequences flanking the TE insertion site are extracted and aligned. However, to handle the LR error rate, the algorithm can accept mismatches (default 1). If the putative TSDs are located more than 2pb away from the detected 5’ and 3’ ends of the TE insertions, they are filtered out because it could be a simple duplication present in the genome. In addition, for OUTSIDER TE insertions, if the putative TSD sequence is not found as a unique sequence at the putative insertion locus in the assembled genome, it is also filtered out and not considered as the sequence of the empty site. Finally, retained TSDs allow to better define the 5’ and 3’ ends of the inserted TE.

(b) To compute the insertion frequency, the number of reads with the TE insertion is divided by the total number of reads spanning the same position, either with or without TE.

#### OUTPUT data

Several BED files are produced containing information on the inserted or deleted TE insertions, their nature and their location on the reference and assembled genomes, as well as the computed frequencies for each of them.

### Using simulated datasets to benchmark the ability of TrEMOLO to annotate new TE insertions on a reference genome

We compared the results obtained by TrEMOLO, TELR (24) and TLDR (25), two recently developed computational tools dedicated to TE detection using LR sequencing data. The rationale of these tools is similar to the one used with short-read sequencing data: they use LR data mapped to a reference sequence, through a BAM file for TLDR (25) or directly from the mapping tool output for TELR. Insertions are detected based on discordant mappings (*i.e*. clipped reads and partially mapped reads). Finally, as for short-read-based approaches, the non-mapped parts of the reads are aligned against a library of canonical TE sequences. Then, discordant mapped reads associated with TE matches are selected, clustered, and analyzed to validate or invalidate the TE insertion sites in the reference genome.

We generated a simulated reads dataset (sample S1, see Methods and Fig. S2) in which 1100 TE insertions with different allele frequencies were added to the *D. melanogaster* unmasked genome called G0-F100 previously published (13). For that, we first generated LR dataset with DeepSimulator on G0-F100 (sb1 at a depth of 10x). Then we used a G0-F100 pseudo genome containing 100 TE insertions to simulate the so-called INSIDER TE sequencing dataset (sb2 at a depth of 45x). To obtain the OUTSIDER TE simulated sequencing dataset (sb3 at a depth of 5x), we launched DeepSimulator on the G0-F100 INSIDER pseudo genome into which we had inserted 1000 additional TEs. Finally, the three LR datasets were mixed together (sb1+sb2+sb3) to obtain the sample S1 (see Methods and Fig. S2; simulated data are available at https://doi.org/10.23708/N447VS).

We mapped this simulated reads dataset to the unmasked G0-F100 genome using TELR (24), TLDR (25) and the TrEMOLO OUTSIDER module (here called “TrEMOLO_NA”, see Methods) with minimap2 (30). We also launched the whole TrEMOLO pipeline (both OUTSIDER and INSIDER modules) with the G0-F100 genome as reference and as the assembled genome, we used the G0-F100 genome with the 100 highly frequent TE insertions. As the TLDR parameters are fixed (25), we decided to use the same parameters for the four methods/tools (TrEMOLO, TrEMOLO_NA, TLDR, and TELR): 200 bp as minimum TE insertion length and 80% as minimum TE sequence identity. To limit false-positive TE detections, we also set at 15kb the largest TE insertion length.

Using these simulated datasets, we observed that out of the 1100 simulated insertions, TELR and TLDR only detected 12 (1%) and 928 (84.4%) TE true-positive insertions, respectively. By contrast, TrEMOLO_NA found 1064 (96.7%) TE true-positive insertions, illustrating the greater sensitivity of the TrEMOLO mapping algorithm (Fig. 2). Unlike TLDR and TELR tools, TrEMOLO also relies on LR based genome assemblies, which enables it to retrieve and use the information of the most frequent TE polymorphims. When using a complete TrEMOLO approach, *i.e*. by taking advantage of the comparison of two whole-genome assemblies, the detection rate even rose up to 97.3%.

**Figure 2:**
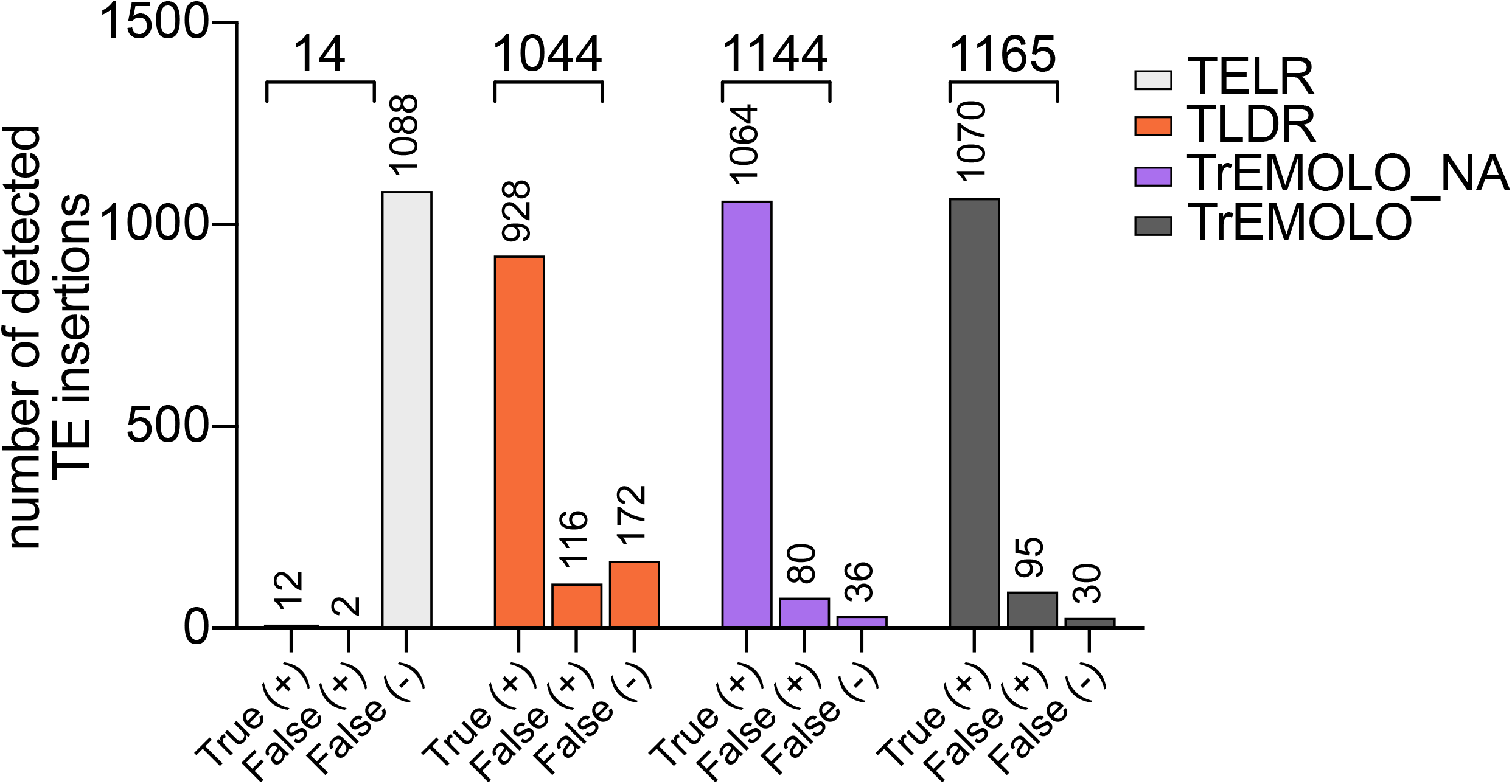
TrEMOLO benchmarking using simulated LR sequencing data. The number of truepositive (True (+)), false-positive (False (+)) and false-negative (False (-)) results obtained using the simulated data (1100 known TE insertions) and four methods/tools (TELR, TLDR, TrEMOLO_NA and TrEMOLO).

Moreover, only 80 (7%) and 95 (8%) TE false-positive insertions were detected in TrEMOLO_NA and TrEMOLO respectively, which is the same order of magnitude as the TLDR false positive insertion rate (11%), and remains very low compared to the significant gain of true-positive insertions detected by TrEMOLO.

### TrEMOLO experimental validation

To validate TrEMOLO insertion/deletion detection accuracy with real biological samples, we took advantage of a previously described *D. melanogaster* laboratory line (36) in which the knockdown of Piwi, a protein essential for TE silencing, can be induced in adult follicle cells (somatic cells of the ovary). This transient somatic Piwi knockdown allows TE derepression in follicle cells and their integration as new proviruses in the progeny genome (36). We performed TE derepression for 73 successive generations to obtain the G73 line that contains thousands of newly integrated TEs in its genome (13,36). We used genomic DNA extracted from G0-F100 (before Piwi knockdown) and G73 (G73-SRE; see Methods) fly samples to check by genomic PCR, 12 of the 2334 newly integrated TEs detected by TrEMOLO in a previous G73 sequencing dataset (G73 depth 183x) (13).This small sample is representative of both OUTSIDER (mostly from the gtwin and ZAM families) and INSIDER TE insertions, i.e. insertions detected by TrEMOLO in either a minority or a majority of the initial G73 (183x) LRs, respectively. Using G73-SRE DNA, genomic PCR yielded amplicons of the expected sizes, whereas no amplification was observed when the G0-F100 reference genomic DNA was used as a template (Fig. 3a, upper panels and lower left panel). Moreover, to control the DNA sample quality, we also checked the presence of four TEs shared by both genomes (Fig. 3a, lower right panel). We also experimentally validated the excision of a *hobo* DNA transposon in G73 that was present in the G0-F100 genome (Fig. 3b; 3c). Altogether, these data illustrated the efficacy of TrEMOLO to detect both insertions and excisions whatever their frequency in the population.

**Figure 3:**
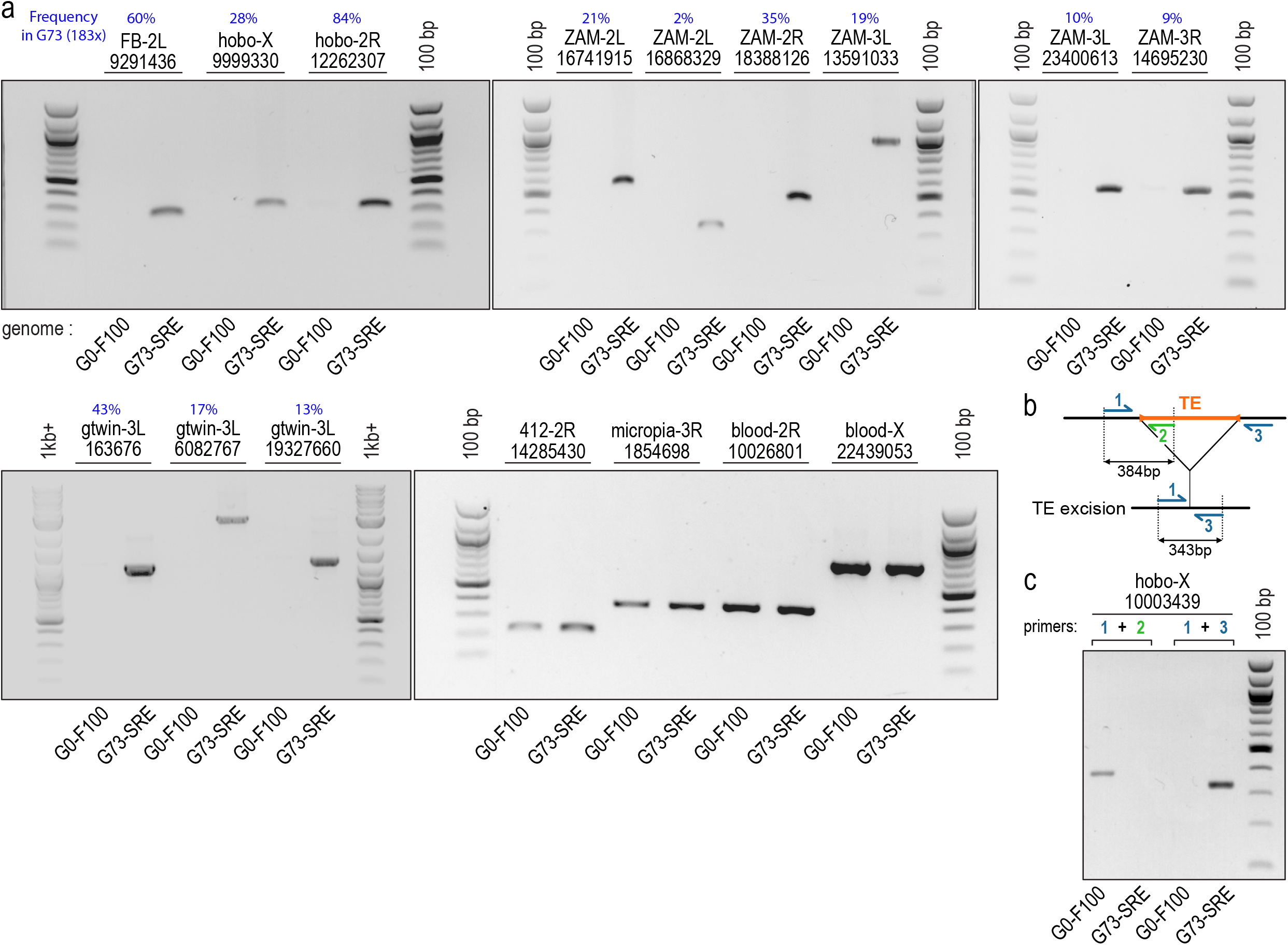
Experimental validation of TrEMOLO specificity of TE insertion/deletion detection. a) PCR performed using long DNA molecules extracted from the G0-F100 (control) and G73 Drosophila lines (genomic DNA extracted using Short Read Eliminator kit: G73-SRE sample). The name of the TE family identified by TrEMOLO is indicated followed by the chromosome arm and the genomic coordinate in the G73 (183x) assembly. The frequencies determined in the G73 (depth 183x) of each TE insertion tested by PCR are indicated as percentages in blue. b) Illustration of the primers used to detect a hobo excision c) PCR performed using long DNA molecules extracted from the G0-F100 (control) and G73 Drosophila lines. The primer sequences and amplicon expected sizes are in Table S1.

### Impact of the genome assembly quality on TrEMOLO efficiency

To assess the impact of the genome assembly quality on TE insertion/deletion detection, we ran TrEMOLO using the G73 genome assembled with different tools and after different polishing rounds, using the G0-F100 assembly as reference. We used Flye (37), Shasta (38) and Wtdbg2 (39) followed by Racon (40) polishing steps and a scaffolding by RaGOO (41) (Methods). As previously reported (13), four rounds of Racon polishing (40) increased the BUSCO score (Table 1), suggesting a better-quality assembly.

**Table 1 :** Comparison of the TE insertion/deletion detection efficiency by TrEMOLO using genome assemblies of different quality.

| Assembly process | Contigs / scaffolds number | N50 (bp) | BUSCO | Assembly total size (bp) | Number of TE insertions | Number of INSIDER TE insertions | Number of OUTSIDER TE insertions | Number of INSIDER TE deletions |
| --- | --- | --- | --- | --- | --- | --- | --- | --- |
| Flye + Racon (4 runs) + Ragoo* | 30 | 27 282 734 | 98% | 148 869 489 | 2334 | 60 | 2274 | 49 |
| Flye + Racon (1 run) + Ragoo* | 31 | 27 316 827 | 96.9% | 149 145 078 | 2339 | 54 | 2285 | 49 |
| Shasta + Ragoo* | 15 | 27 256 021 | 93% | 151 203 880 | 2344 | 94 | 2250 | 56 |
| Wtdbg2 + Ragoo* | 22 | 34 296 233 | 92.6% | 171 989 387 | 2579 | 159 | 2420 | 109 |

\* Ragoo was run with default parameters.

Using genomes assembled with the recently updated Flye and Shasta assemblers led to the detection of similar numbers of INSIDER and OUTSIDER TEs, with a slightly higher number of INSIDER TEs for the Shasta-assembled genome (Table 1). When using a genome assembly generated with Wtdbg2, which has not been updated for two years and thus has not incorporated the recent changes in ONT sequencing data (*e.g*., new base calling inducing a much lower error rate) and has a lower BUSCO score (indicating a lower quality of assembly), TrEMOLO detected roughly the same number of OUTSIDER TE insertions as the more recent assemblers. Note however, that the INSIDER module produced at least twice as much insertions and deletions (among which many of them were probably false positives) probably because it was based on a low-quality assembly.

Altogether, we conclude that a draft genome assembly, even obtained with a not-so-recent assembler version (with a lower BUSCO score indicating a lower assembly quality), is of enough quality however to allow the identification of TE polymorphisms by TrEMOLO.

### Validation of TE frequency estimated by TrEMOLO using simulated data

To validate the accuracy of TrEMOLO frequency estimation, we used simulated datasets with known frequencies of INSIDER and OUTSIDER TEs (see Methods; Fig. 4a). The frequency is computed by the number of reads supporting the insertion divided by the total number of reads spanning the region of insertion. For OUTSIDER TEs, we count as supporting reads, the reads with the complete insertion as well as the clipped ones (Fig. 4a). TrEMOLO frequency module slightly overestimated the frequency of INSIDER TEs for the simulated datasets (Fig. 4b). For OUTSIDER TEs, whatever is the frequency, TrEMOLO obtained an accurate estimation of the frequency (Fig. 4c).

**Figure 4:**
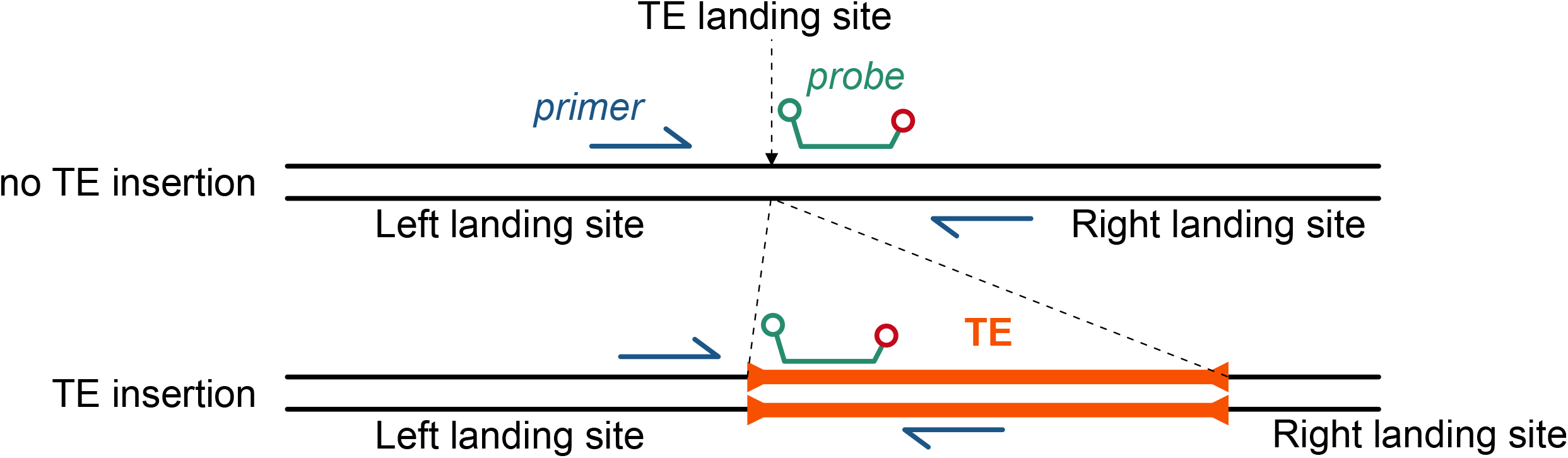
Validation of TrEMOLO frequency estimation. a) Rationale of frequency estimation using fully mapped and clipped reads. b) Violin plots of the INSIDER TE insertion frequencies estimated by TrEMOLO using the simulated datasets. Dashed lines represent the real frequency in the simulated datasets. c) Violin plots of the OUTSIDER TE insertion frequency estimated by TrEMOLO in the simulated datasets with clipped reads. Dashed lines represent the real frequency in the simulated datasets.

### Experimental validation of TE frequency estimated by TrEMOLO using droplet digital polymerase chain reaction

To validate the TrEMOLO frequency module accuracy, we re-extracted long DNA molecules from G73 flies (Methods) and analyzed the extracted DNA by ONT sequencing with Short-Reads Eliminator kit (G73-SRE library) and by droplet digital polymerase chain reaction (ddPCR). This latter approach allows the absolute quantification using forward primers located in the left side of the TE landing site and reverse primers located either in the TE or on the right side of the landing site (Fig. 5). To increase the detection specificity, we also designed fluorescent probes. The primer and probe sequences are listed in Table S2. We selected nine of the 2234 TE insertions, as representatives of different TEs families and of frequencies in the initial G73 library (depth of 183x) (13). We ran TrEMOLO (parameter: 80% as minimum TE sequence identity and 10% as minimum TE insertion length) and estimated their frequency in the new G73-SRE library having a depth of 82x (Table 2).

**Figure 5.**
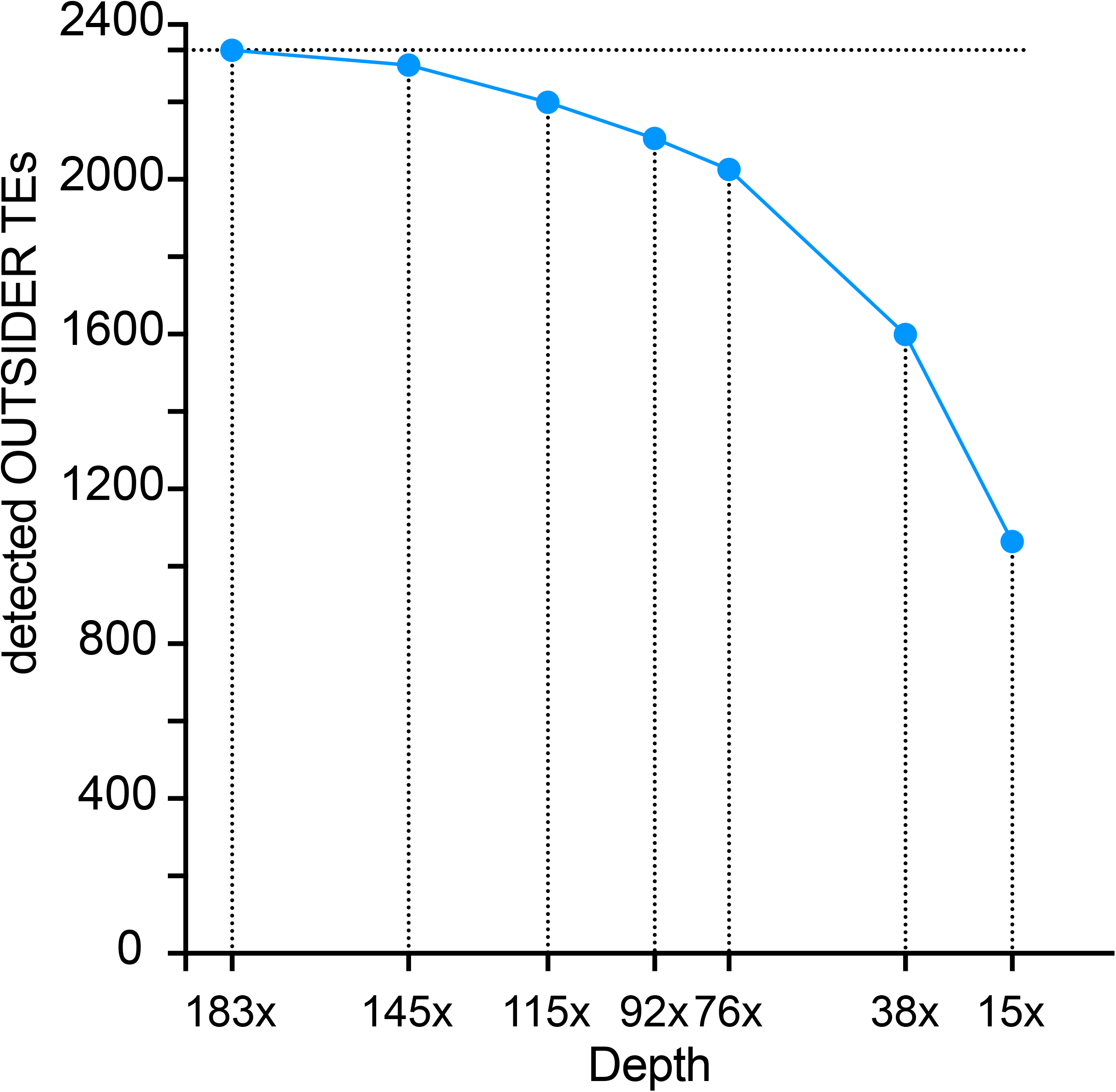
Example of primer pair and probe design for ddPCR quantification of the “empty” (no TE insertion) or the “full” (TE insertion) haplotype. Primers and probes for each tested TE are listed in Table S2.

**Table 2 :** TE frequencies estimated using ddPCR and TrEMOLO.

| Name of the TE<br>and<br>its coordinates on<br>G73 (183x) assembly | Frequency estimation (%) |  |
| --- | --- | --- |
|  | ddPCR<br>G73-SRE | TrEMOLO<br>G73-SRE (82x) |
| ZAM-3L_13591033 | 0.6 | NA |
| blood-2L_18128567 | 2.2 | 37.68 |
| ZAM-2L_16741915 | 9.6 | 17.3 |
| gtwin-3L_6082767 | 10.4 | 6.58 |
| ZAM-2L_16868329 | 13.6 | 17.89 |
| ZAM-2R_18388126 | 17.3 | 18.31 |
| gtwin-3L_163676 | 26.5 | 27.63 |
| roo-2L_13882749 | 59.2 | 57.95 |
| 412-2R_14285430 | 82.6 | 93.44 |

NA= TE insertions with no reads containing the insertion in G73-SRE (82x).

For two of them, the low frequencies estimated by ddPCR (0.6% and 2.2%) do not allow an accurate estimation of their frequency by TrEMOLO. In the G73-SRE library (depth 82x) there are no reads supporting the ZAM-3L_13591033 insertion, while we detected it using the initial library G73 that has shorter reads and a higher depth (183x). Concerning the blood-2L_18128567 insertion, the TrEMOLO frequency estimation is not congruent with the ddPCR one. This overestimation of the frequency by TrEMOLO (37.68%) illustrates that TrEMOLO estimation starts to be accurate when the insertion TE frequency is around 10% in the population. Indeed, for the other six TE insertions tested, the ddPCR estimates fitted very well with the TrEMOLO estimates, whatever their frequency from 10% to 82%.

### Impact of sequencing depth on TE calling and frequency estimation

The total amount of the initial G73 sequencing is averaging a depth of 183x, fairly representative most of the 200 haploid genomes of the 100 males used for the DNA extraction. Indeed, the probability of sequencing one out of the 200 haplotypes from this sample (i.e. the depth per haplotype) was (1:200)x183=0.91. From this dataset, we generated several sequence subsets using Filtlong (42) (see Methods), resulting in seven subsamples of reads (ranging from 145x to 15x of depth) (Table S3). We then launched TrEMOLO with these datasets, and compared the number of OUTSIDER TE insertions detected by mapping these reads to the G73 assembly (Table S3 and Fig. 6).

**Figure 6.**
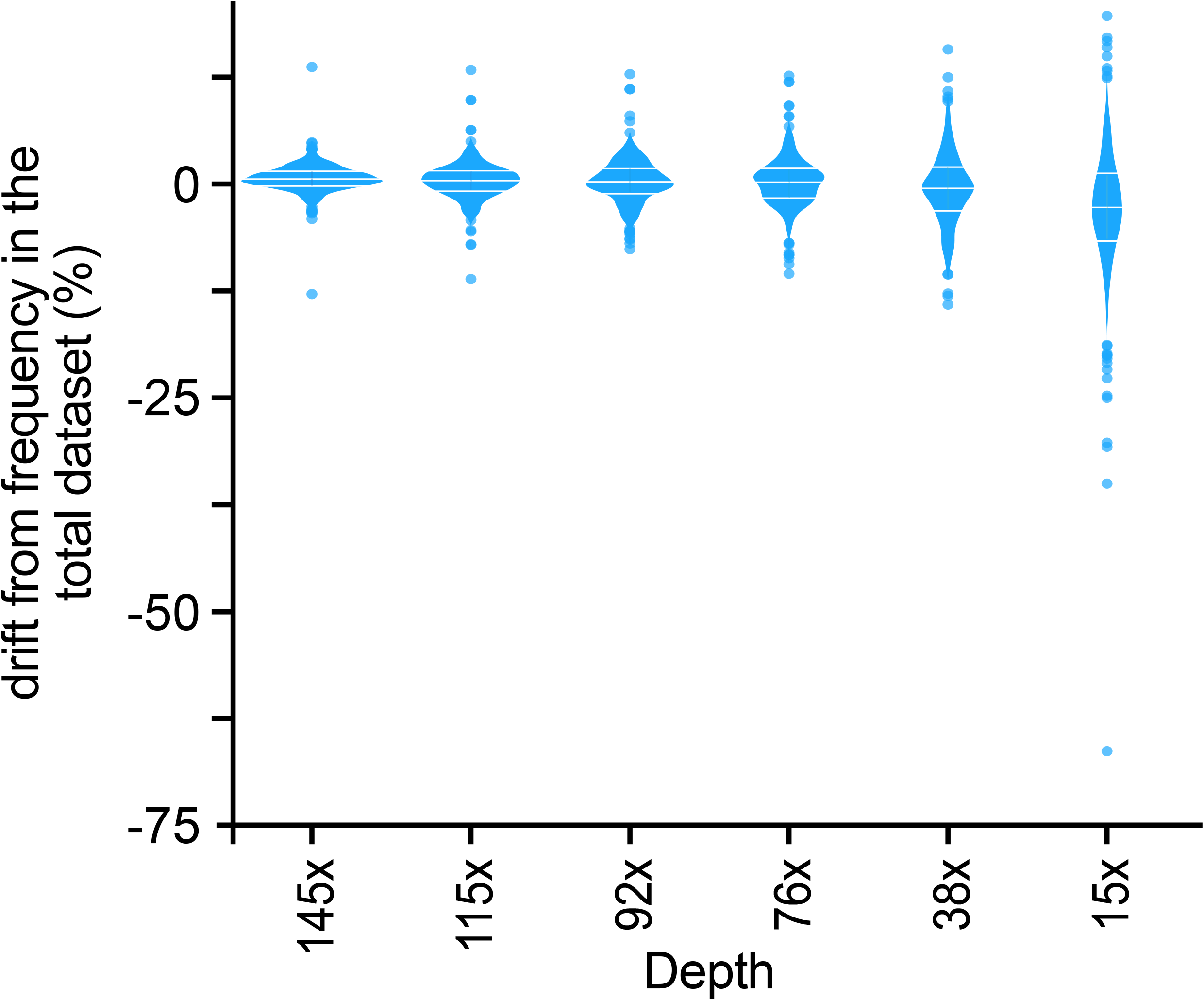
OUTSIDER TEs detection depends on depth of sequencing. G73 sequencing reads averaging a 183x depth were subsampled and the OUTSIDER TrEMOLO module was run to detect OUTSIDER TE insertions at each depth.

Using the initial dataset G73 (183x), TrEMOLO detected 2334 TEs of which 2274 are OUTSIDER TE insertions. The percentage of detected OUTSIDERs is relative to this dataset. Decreasing the depth from 183x to 76x slightly reduced the number of OUTSIDER TEs detected. Then, TrEMOLO detection performance decreased more rapidly. Nevertheless, even with the lowest depth (15x), TrEMOLO could still identify 47% (1064) of OUTSIDER TE insertions. As expected, these insertions were significantly more frequent in the 183x dataset than the 1210 ones that were lost during the down-sampling (Fig. S3, *p*=0,05 Wilcoxon rank test). Thus, in case of highly polymorphic populations, such as ours, we would recommend a sequencing depth corresponding to a minimum of 0.7x per haplotype (corresponding here to our 145x dataset) to ensure an optimal recovery of low-frequency transpositions.

Among the 2334 original TE insertions, only 820 OUTSIDER TEs were detected in all subsamples (Fig. 7). To test the impact of the sequencing depth on the accuracy of frequency estimates, we computed the frequencies of these 820 shared TE insertions in each subsample. The accuracy of OUTSIDER TE insertion frequency estimation was acceptable when using datasets with sequencing depth ≥76x, with a decrease of 12% maximum compared with the initial frequency computed at 183x (Fig. 7). However, for very “low depth” datasets, (here 0.19x per haplotype or less), the accuracy frequency decreased quite rapidly with an almost systematic underestimation by up to 70% compared with the original dataset.

**Figure 7.**
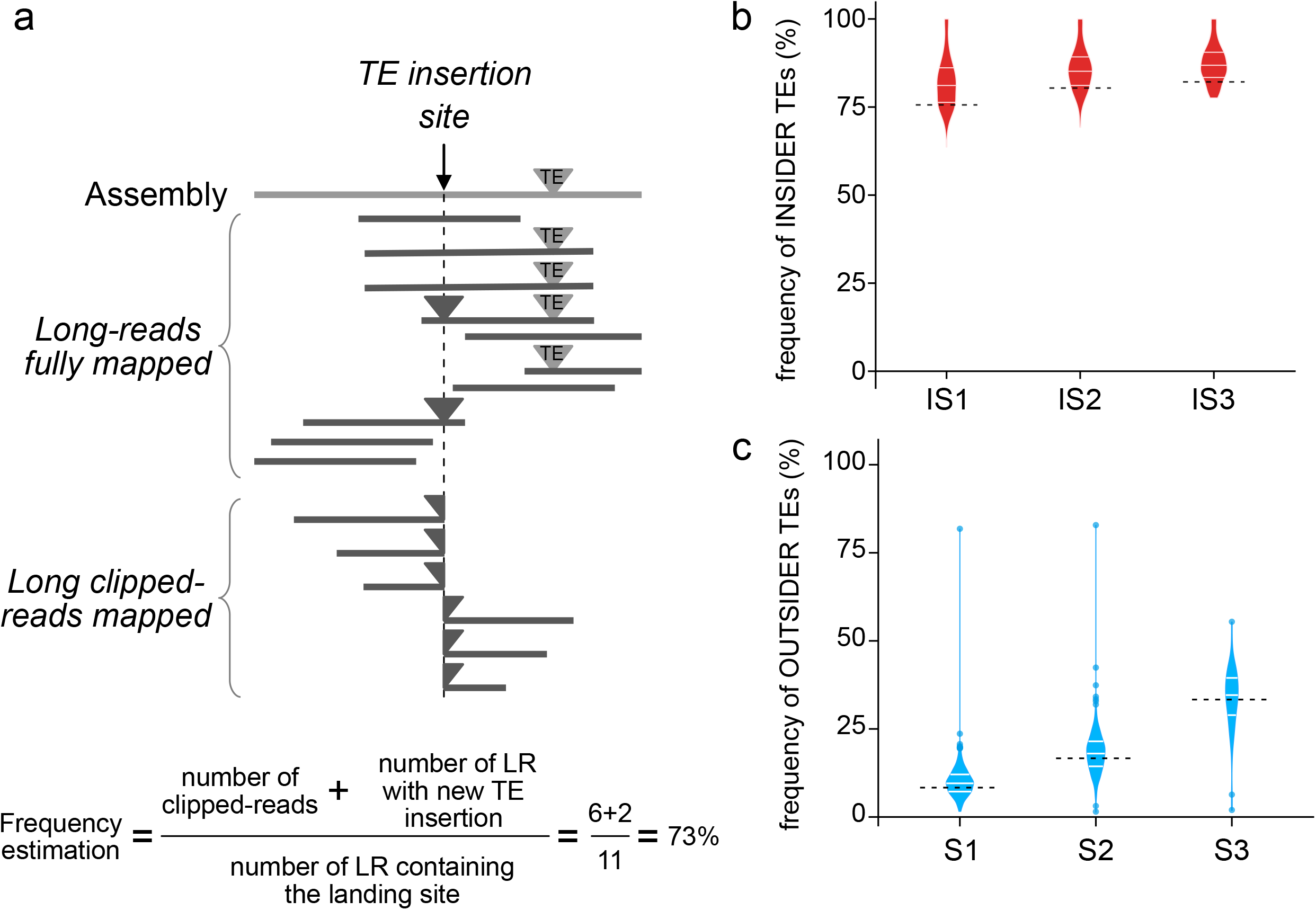
TrEMOLO frequency estimation variation of the 1064 common OUTSIDER TEs using datasets of different depths. Violin plot of the difference (positive or negative ratio) between the frequency computed for each common insertion in a given dataset (different sequencing depths) and the frequency of the same OUTSIDER TE insertion computed in the total dataset.

## Discussion

Accurate detection of TEs with different frequencies in a population or in a single sample at the somatic level is important for researchers in evolution, ecology, agronomy, and medicine. However, it is still a challenge due to the repetitive nature of the TEs. Most of the current approaches to identify TE insertions are based on short-read mapping to a reference genome, sometimes quite distant from the studied samples. Working with distantly related genomes may drastically bias the mapping results and thus the subsequent analyses, for instance by providing high false positive rates (43). Therefore, TE detection tools based on mapping approaches allow the identification of low-frequency TE insertions (or from the non-major haplotype), but not all of them, only when using very high-depth short-read sequencing data. The availability of low-cost LR sequencing technologies (ONT) opens the possibility to accurately detect new TE insertions. Improving the sequencing quality allows to obtain accurate and (almost) complete genome assemblies in a short amount of time for small sized ones. For instance, a high-quality *Drosophila* genome can be assembled in 24 hours (44). However, most of these assemblers provide a single haplotype sequence as output, generally the most prevalent. Finally, most TE detection tools rely on the identification of TE fragments as source of SV *ab initio*, through a global genome annotation without any comparison with other already annotated sequence.

We designed TrEMOLO, a comprehensive computational tool with the aim of combining the main haplotype information (assembly step) followed by the recovery of minor haplotypes by LR sequence mapping to the newly assembled genome to detect all SV types without *ab initio* TE detection. It is only after this naive detection that the SVs are annotated as possible TEs or not. This approach allows to run TrEMOLO using different TE databases. TrEMOLO accuracy in TE identification and the correction system used for TSD detection allow predicting the insertion site within a 2-base pair window (provided, of course, that the extremities of the TE family are precisely defined in the consensus sequence). Therefore, TrEMOLO is currently one of the most efficient tools for TE detection. Moreover, TrEMOLO allows the estimation of the insertion frequency for each detected TE copy with high accuracy (when using data with sequencing depth >50x). Importantly, TrEMOLO performance is weakly influenced by the assembly quality and sequencing depth variations (up to a certain point). In terms of parameters, TrEMOLO in its standard approach will use the previously published 80/80/80 rule for TE recovery (1). Indeed, it relies on 80% of similarity to and 80% in length of the reference TE sequence. However, as shown in the tests before, we could decrease the length coverage down to 10% and still recover true positive variations. This length parameter can thus be modified for e.g. identifying partially transposed elements (for insertions of 5’-truncated LINEs), or insertions of abnormal copies (internally deleted TEs still able to transpose). In the same way, decreasing the level of similarity would allow to detect more distantly related copies from the consensus/reference copy in the database. However, in this case, the decrease of the stringency of detection will have a stronger effect on the quality of results (improving the false positive rates) (45), and we do not recommend it. The best approach would be to add in the reference TE file the targeted sequences instead.

In terms of execution times and memory usage, it depends a lot on the considered species. However, as an example, the analyses presented here for simulated data were run on a single laptop computer with 16 Gb of RAM and an 12-cores i7 Intel processor, in less than 10 hours, and thus can be deployed in almost any lab.

Therefore, TrEMOLO can be used for different projects (e.g. pooled DNA sequencing, population analysis, human genetics, tumor analysis) that require the fast and precise identification of TE insertions, even at very low frequency within a sample. Indeed, our data showed that a single insertion can be unambiguously identified through a unique read, as long as the read is long enough to harbor the whole TE and its surrounding insertion site.

## Conclusions

TrEMOLO allows the precise analysis and study of haplotype-specific TE insertions and a better understanding of the true impact of TE evolution in the wider context of their host genome.

## Methods

### TrEMOLO pipeline preprocessing

INSIDER TE identification: to compare the reference genome and a new assembly, TrEMOLO uses the same methodology as RaGOO (v1.1) (41) up to the SV identification step. Then, SVs are compared to the TE database (Bergman’s laboratory, https://github.com/bergmanlab/drosophila-transposons/) with BLASTN (2.9.0) (--outfmt 6) (46). The output file is filtered based on the size and identity percentages listed in parameters (80% in standard for each of them). DNA fragments that appear to be part of the same sequence are grouped. For INSERTION, Repeat_expansion, Tandem_expansion, sequences are retrieved from the query (new assembly). For DELETION, Repeat_contraction, Tandem_contraction, sequences are recovered from the reference (Fig. S1)

OUTSIDER TE identification: the LRs are aligned on the genome resulting from their assembly using minimap2 (v.2.24) (30) and SVs are identified using Sniffles (47). Reads covering the SV are recovered from the SAM/BAM file using as anchor point the Sniffles provided position and a 30bp-window with SAMtools (48). The SV reads and sequences are compared to the TE database, and filtered as done for INSIDER TEs, but only the best support reads are kept among those provided by Sniffles or the SAM parsing (Fig. S1).

TSD detection: both TE flanking sequences are recovered to check whether there are duplicate sequences at the TE junction (see Results for more details). For an OUTSIDER TE, if TrEMOLO detects duplicated sequences farther than the initially identified TE junctions, and if they fit to the TSD, then the OUTSIDER TE position is corrected accordingly.

TE frequency estimation: two different approaches are used to estimate the INSIDER and OUTSIDER TE frequencies in the samples. For INSIDER TEs, from the original BAM file only reads aligned to the putative TE positions (with a 100bp window 5’ and 3’) are retained using SAMtools view and the -F 2048 -L option (removing any reads with these FLAGS value (see SAM specifications) and overlapping a provided BED file, respectively). Then, clipped reads are removed to keep only the reads aligned on the putative TE position within a 5bp window (5’ and 3’). The mean depth of 30bp at the 5’ and 3’ of the putative insertion is computed with SAMtools depth. Then, the TE frequency is computed as (number of reads showing the insertion)/(total number of reads).

For OUTSIDER TEs, first, reads not aligned to the putative TE insertion site are removed from the original BAM file (SAMtools view with the -l option). Then, the clipping tags in the CIGAR strings for the remaining reads are modified by adding an insertion tag of the TE size plus 10 bp to the read sequence (*i.e*., the same number of “N” at the corresponding position with a fake quality of 8). Then, Sniffles is run again to obtain the true number of reads associated with a given TE. After recovering the local depth, as done for INSIDER TEs, frequency is computed as (number of reads showing the insertion)/(total number of reads).

### Simulated datasets

A previously assembled reference *D. melanogaster* genome, called G0-F100 (13) was used for the LR simulation by DeepSimulator (version v1.5) (49) (-B 2 -K 10 -l 20 000) with a mean length of 20kb and a depth of 10X (subsample 1 (sb1)) (Fig. S2). In parallel, two fake genomes containing INSIDER TE insertions only or both INSIDER and OUTSIDER TE insertions were built as follows. For INSIDER TE insertions, 100 TE sequences were artificially and randomly inserted in the G0-F100 assembly genome. They correspond to the canonical sequences of 11 TE families of different classes and different lengths (from one to 13kb), each one providing five (ZAM and gtwin) or ten new insertions (Doc, roo, copia, F-element, hopper, Helena, HB, TART-A and gypsy). Then, DeepSimulator (version v1.5) (47) (-B 2 -K n -l 20000) was used to simulate LRs with a mean length of 20kb and depth of 30, 40 and 45x (subsamples 2 (sb2)). For OUTSIDER TE insertions, 1,000 TE sequences belonging to the same families as the simulated INSIDER TEs and also to the Idefix family (200, 200, 50, 50, 50, 50, 50, 50, 50, 50, 100, and 100 insertions, respectively) were inserted randomly in the G0-F100 genome assembly containing the INSIDER TE insertions. Then, DeepSimulator (version v1.5) (49) (-B 2 -K n -l 20000) was used to simulate LRs with a mean length of 20kb mean length and depth of 5, 10 and 20X (subsamples 3 (sb3)).

These three LR subsamples were mixed together (total depth of 60X) to test TrEMOLO sensitivity at different levels: S1: (10X sb1 +45X sb2 + 5X sb3), S2: (10X sb1 + 40X sb2 + 10X sb3), and S3: (10X sb1 + 30X sb2 + 20X sb3) (Fig. S2).

For the INSIDER TE frequency estimation, the sb1 and sb2 subsamples were used together with a new sb4 subsample simulated with a depth of 1X and minimum to maximum length of 10 to 50kb to obtain IS1 (10X sb1 + 30X sb2 + 1X sb4), IS2 (10X sb1 + 40X sb2 + 1X sb4) and IS3 (10X sb1 + 45X sb2 + 1X sb4). Simulated data set are available at https://doi.org/10.23708/N447VS.

### Tools comparison

For TLDR (v1 - February 2021; (25)), the specified parameters were: --min_te_len 200 -- max_te_len 15000 (minimum of 200 nt and maximum of 15 000 nt in size for detected TEs). The same parameters were used for TrEMOLO and TrEMOLO_NA. For TELR (v1 - August 2021; (24)), the settings were --minimap2_family --assembler flye -x ont --aligner minimap2.

### Genome assembly

Raw ONT reads for the G73-SRE sample were QC checked using Nanoplot v1.10.2 (https://github.com/wdecoster/NanoPlot) for sequencing run statistics. Reads with QC <7 were removed by the sequence provider (Montpellier Genomix, Montpellier, France) before QC. For all ONT sequencing data, the genome was assembled as previously published (13), except that a newer version of Flye (v2.9) was used.

### Datasets

The LR dataset of the *D. melanogaster* laboratory G0-F100 and G73 lines previously published (13) is available at ENA (https://www.ebi.ac.uk/ena) under the accession number ERP122844. Simulated data set are available at https://doi.org/10.23708/N447VS.

### Benchmark of *Drosophila* genome assemblies effect

Genome assemblies were performed using the raw LR data and CulebrONT for assembly and polishing (50). Flye v2.9 (37) was used with standard parameters except for the --nano-raw option. Four rounds of Racon v1.4.3 with minimap2 v.2.24 (30) were performed using the standard conditions for ONT data, and with medaka v1.2 (standard conditions). Scaffolding was performed using RaGOO v1.1 (41) with standard options except for output structural variations with the -s option and the corresponding reference. For raw assemblies, Wtdbg2 v2.5 (39) and Shasta v0.1.0 (38) were used (standard conditions).

### Impact of sequencing depth on TE calling and frequency estimation

A previously published G73 long read sequencing library (13) was downsized to smaller samples using FiltLong (42) and the following options: -t 19000000000, -t 15000000000, -t 12000000000, -t 10000000000, -t 5000000000, -t 2000000000). This subsampling resulted in a series of sequencing datasets with more or less degraded depths ranging from 183x to 15x.

### Genomic DNA extraction and Short-Read Eliminator kit (SRE)

Genomic DNA was extracted from 50 mg of G0-F100 and G73 male flies using the Nanobind Tissue Big DNA Kit (Circulomics SKU NB-900-701-01) with Insect buffer (PL1) as lysis buffer. Briefly, samples were homogenized in 200 μl CT buffer with a pellet pestle, centrifuged at 16,000 x g at 4°C for 2min. Pellets were washed in 1 ml CT buffer and centrifuged at 16,000 x g at 4°C for 2min. Pellets were resuspended in 200μl PL1 lysis buffer (Circulomics Aux. Kit) supplemented with 20μl proteinase K and incubated in a thermomixer at 55°C and 900 rpm for 1 h. 20μl RNase A was added and incubation was continued for 15 min. Samples were centrifuged at 16,000x g for 5 min, and supernatants passed through a 70μm filter and collected in a 1.5 ml DNA LoBind microcentrifuge tube. 50 μl BL3, a Nanobind disk and 400 μl isopropanol were added to the samples. Incubation and washes were done according to the manufacturers’ instructions. Elution was performed with 50 μl EB buffer. The genomic DNA concentration was estimated with a Nanodrop spectrophotometer (Thermo Fisher Scientific) and a Qubit Fluorometer with the Qubit dsDNA BR Assay Kit (Thermo Fisher Scientific), according to the manufacturer’s instructions. To get rid of reads <25 kb, genomic DNA samples with concentration adjusted between 50-150 ng/μl in 60 μl were treated with the SRE Kit, 25 kb cutoff (Circulomics SKU SS-100-101-01), according to the manufacturer’s instructions, and eluted in 50 μl EB.

### Library preparation for G73-SRE LR sequencing (depth 82x)

Libraries were prepared on 1μg genomic SRE DNA with the SQK-LSK110 Kit (Nanopore) following the manufacturer’s instructions. 12 μl of each library, 37.5μl of sequencing buffer II (SBII), and 25.5 μl of loading beads II (LBII) were loaded onto the R9.4.1 flow cell. Base-calling was performed using the Guppy v5 hac model.

### PCR amplification of long DNA molecules

PCR was performed with 60 ng G0-F100 or G73 genomic DNA in a final volume of 20μl containing 10μl Platinum Hot-Start Green Master Mix 2X (Invitrogen 14001-012) and 0.2 μM final concentration of each primer (see Table S1 for sequences). Annealing was performed at 60°C for 15 sec and elongation was adapted to the amplicon size (15 sec/kb). Between 3 and 5 μl of each PCR reaction were loaded on a 1% agarose gel with EtBr.

### Droplet digital PCR for TE insertion frequency determination

Between 5 and 15 ng genomic SRE DNA was combined with 10 μl of 2x ddPCR Supermix for probes (No dUTP) (Bio-Rad), 1μl 20x primers/probe sets (1x primers/probe set includes 450 nM each primer and 250 nM probe) for the targets (see design in Table S2), 1μl 20x primers/probe set for the housekeeping reference gene (*RpL32*), 1μl restriction enzyme (*HindIII*, diluted 1:10 in diluent B (NEB)) and nuclease-free water to a total volume of 20μl.

For droplet generation, 20 μl of each mixture was transferred to the center line of a DG8 droplet generator cartridge (Bio-Rad Laboratories). Then, 70 μl of droplet generation oil for the probe (Bio-Rad Laboratories) was injected in the cartridge bottom line and droplets were produced using a QX200 Droplet Generator (Bio-Rad Laboratories). Droplets (40 μl) were transferred to a 96-well plate and endpoint PCR was performed on a T100 Thermal Cycler (Bio-Rad Laboratories) using the following cycling protocol: enzyme activation at 95 °C for 10 min, followed by 40 cycles of 95 °C for 30 s and 60 °C for 1 min (ramp rate settings of 2°C/sec), followed by a droplet stabilization step at 98 °C for 10 min, and a final hold step at 4 °C. Droplet measurement was performed on a QX200 Droplet Reader (Bio-Rad Laboratories) and data analyzed with the QuantaSoft software (Bio-Rad Laboratories). Two different fluorescent signals were measured: i) amplification of the TE inserted in a given locus sequence (*TE_in_*) or amplification of the same locus without TE (*TE_out_*) in the FAM channel, and ii) amplification of the *RpL32* housekeeping gene in the HEX channel. The proportion of *TE_in_* or *TE_out_* to *Rpl32* was calculated as follows: *TE_in_/Rpl32 or TE_out_/Rpl32*. Each value was expressed as copies/μl. Frequency was calculated with the following formula: Frequency in %= (*TE_in_/Rpl32*x100) / (*TE_in_/Rpl32* + *TE_out_/Rpl3*).

## Declarations

### Availability of data and materials

The LR sequencing data used for this study were deposited at ENA (https://www.ebi.ac.uk/ena) under the accession numbers (ERP138838 under validation by ENA) and *the benchmarking datasets and the command lines are available at https://dataverse.ird.fr/dataverse/tremolo_data*.

- Project name: TrEMOLO
- Project home page: https://github.com/DrosophilaGenomeEvolution/TrEMOLO
- Archived version: https://doi.org/10.23708/2FYBUL
- Operating system(s): Linux
- Programming language: Python/SnakeMake
- Other requirements: Singularity or minimap2/assemblytics/samtools
- License: GNU GPL v3

Any restrictions to use by non-academics: license acceptation

### Authors’ contributions

F.S., M.M., A.S.F.L and S.C. designed the methodology; M.M. and F.S. encoded software; M.V. tested the software; F.S., B.M. and S.C. supervised M.M., K.A. and B.M. respectively; Biological experiments: B.M and K.A.; F.S., A.P. and S.C. wrote the manuscript. Funding acquisition: A.S.F.L, S.C.. All authors have read and agreed to the published version of the manuscript.

## Acknowledgments

We thank Dr Marie Fablet for her comments on the manuscript; Elisabetta Andermarcher for manuscript editing; We acknowledge the ISO 9001 certified IRD itrop HPC (member of the South Green Platform) at IRD Montpellier for providing HPC resources that have contributed to the research results reported in this paper (URLs: https://bioinfo.ird.fr/ and http://www.southgreen.fr); the Genotoul platform (https://genotoul.fr/) and (https://www.france-bioinformatique.fr/) for providing calculation time on their servers.

## References

1. Wicker T, Sabot F, Hua-Van A, Bennetzen JL, Capy P, Chalhoub B, et al. A unified classification system for eukaryotic transposable elements. Nat Rev Genet. 2007 Dec;8(12):973–82.

2. Jiang N, Feschotte C, Zhang X, Wessler SR. Using rice to understand the origin and amplification of miniature inverted repeat transposable elements (MITEs). Curr Opin Plant Biol. 2004 Apr;7(2): 115–9.

3. Macas J, Neumann P. Ogre elements--a distinct group of plant Ty3/gypsy-like retrotransposons. Gene. 2007 Apr 1;390(1–2): 108–16.

4. Pritham EJ, Putliwala T, Feschotte C. Mavericks, a novel class of giant transposable elements widespread in eukaryotes and related to DNA viruses. Gene. 2007 Apr 1;390(1–2):3–17.

5. Walkowiak S, Gao L, Monat C, Haberer G, Kassa MT, Brinton J, et al. Multiple wheat genomes reveal global variation in modern breeding. Nature. 2020 Dec;588(7837):277–83.

6. Mérel V, Boulesteix M, Fablet M, Vieira C. Transposable elements in Drosophila. In: Mobile DNA. In press.

7. Flutre T, Duprat E, Feuillet C, Quesneville H. Considering Transposable Element Diversification in De Novo Annotation Approaches. PLOS ONE. 2011 Jan 31;6(1):e16526.

8. Flynn JM, Hubley R, Goubert C, Rosen J, Clark AG, Feschotte C, et al. RepeatModeler2: automated genomic discovery of transposable element families [Internet]. Genomics; 2019 Nov [cited 2022 Jul 20]. Available from: http://biorxiv.org/lookup/doi/10.1101/856591

9. Ou S, Su W, Liao Y, Chougule K, Agda JRA, Hellinga AJ, et al. Benchmarking transposable element annotation methods for creation of a streamlined, comprehensive pipeline. Genome Biol. 2019 Dec 16;20(1):275.

10. Roy NS, 최지영, Lee SI, Soo KN. Marker utility of transposable elements for plant genetics, breeding, and ecology: a review. Genes and Genomics. 2015;37(2):141–51.

11. Waugh R, McLean K, Flavell AJ, Pearce SR, Kumar A, Thomas BB, et al. Genetic distribution of Bare-1-like retrotransposable elements in the barley genome revealed by sequence-specific amplification polymorphisms (S-SAP). Mol Gen Genet. 1997 Feb 27;253(6):687–94.

12. Flavell AJ, Knox MR, Pearce SR, Ellis TH. Retrotransposon-based insertion polymorphisms (RBIP) for high throughput marker analysis. Plant J. 1998 Dec;16(5):643–50.

13. Mohamed M, Dang NTM, Ogyama Y, Burlet N, Mugat B, Boulesteix M, et al. A Transposon Story: From TE Content to TE Dynamic Invasion of Drosophila Genomes Using the Single-Molecule Sequencing Technology from Oxford Nanopore. Cells. 2020 Jul 25;9(8):1776.

14. Goerner-Potvin P, Bourque G. Computational tools to unmask transposable elements. Nat Rev Genet. 2018 Sep 19;

15. Bogaerts-Márquez M, Barrón MG, Fiston-Lavier AS, Vendrell-Mir P, Castanera R, Casacuberta JM, et al. T-lex3: an accurate tool to genotype and estimate population frequencies of transposable elements using the latest short-read whole genome sequencing data. Bioinformatics. 2020 Feb 15;36(4):1191–7.

16. Fiston-Lavier AS, Barrón MG, Petrov DA, González J. T-lex2: genotyping, frequency estimation and reannotation of transposable elements using single or pooled next-generation sequencing data. Nucleic Acids Res. 2015 Feb 27;43(4):e22.

17. Fiston-Lavier AS, Carrigan M, Petrov DA, González J. T-lex: a program for fast and accurate assessment of transposable element presence using next-generation sequencing data. Nucleic Acids Research. 2011 Mar;39(6):e36–e36.

18. Nelson MG, Linheiro RS, Bergman CM. McClintock: An Integrated Pipeline for Detecting Transposable Element Insertions in Whole Genome Shotgun Sequencing Data. G3: Genes, Genomes, Genetics. 2017 Jan 1;g3.117.043893.

19. Goubert C, Modolo L, Vieira C, ValienteMoro C, Mavingui P, Boulesteix M. De novo assembly and annotation of the Asian tiger mosquito (Aedes albopictus) repeatome with dnaPipeTE from raw genomic reads and comparative analysis with the yellow fever mosquito (Aedes aegypti). Genome Biol Evol. 2015 Mar 11;7(4):1192–205.

20. Kofler R, Gómez-Sánchez D, Schlötterer C. PoPoolationTE2: comparative population genomics of transposable elements using Pool-Seq. Mol Biol Evol. 2016 Aug 2;msw137.

21. Vendrell-Mir P, Barteri F, Merenciano M, González J, Casacuberta JM, Castanera R. A benchmark of transposon insertion detection tools using real data. Mobile DNA. 2019 Dec;10(1): 1–19.

22. Rech GE, Radío S, Guirao-Rico S, Aguilera L, Horvath V, Green L, et al. Population-scale long-read sequencing uncovers transposable elements associated with gene expression variation and adaptive signatures in Drosophila. Nat Commun. 2022 Apr 12;13(1):1948.

23. Disdero E, Filée J. LoRTE: Detecting transposon-induced genomic variants using low coverage PacBio long read sequences. Mob DNA. 2017;8:5.

24. Han S, Dias GB, Basting PJ, Viswanatha R, Perrimon N, Bergman CM. Local assembly of long reads enables phylogenomics of transposable elements in a polyploid cell line. Nucleic Acids Research. 2022 Sep 26;gkac794.

25. Ewing AD, Smits N, Sanchez-Luque FJ, Faivre J, Brennan PM, Richardson SR, et al. Nanopore Sequencing Enables Comprehensive Transposable Element Epigenomic Profiling. Molecular Cell. 2020 Nov;S1097276520307310.

26. Lin J, Jia P, Wang S, Ye K. Comparison and benchmark of long-read based structural variant detection strategies [Internet]. Bioinformatics; 2022 Aug [cited 2022 Dec 8]. Available from: http://biorxiv.org/lookup/doi/10.1101/2022.08.09.503274

27. He K, Minias P, Dunn PO. Long-Read Genome Assemblies Reveal Extraordinary Variation in the Number and Structure of MHC Loci in Birds. Genome Biol Evol. 2021 Feb 3;13(2):evaa270.

28. Nurk S, Koren S, Rhie A, Rautiainen M, Bzikadze AV, Mikheenko A, et al. The complete sequence of a human genome. Science. 2022 Apr;376(6588):44–53.

29. Kurtz S, Phillippy A, Delcher AL, Smoot M, Shumway M, Antonescu C, et al. Versatile and open software for comparing large genomes. Genome Biology. 2004 Jan 30;5(2):R12.

30. Li H. New strategies to improve minimap2 alignment accuracy. Alkan C, editor. Bioinformatics. 2021 Dec 7;37(23):4572–4.

31. Welcome to Python.org [Internet]. Python.org. [cited 2022 Jul 20]. Available from: https://www.python.org/

32. Mölder F, Jablonski KP, Letcher B, Hall MB, Tomkins-Tinch CH, Sochat V, et al. Sustainable data analysis with Snakemake. F1000Res. 2021;10:33.

33. Hedges DJ, Deininger PL. Inviting instability: Transposable elements, double-strand breaks, and the maintenance of genome integrity. Mutat Res. 2007 Mar 1;616(1–2):46–59.

34. Nattestad M, Schatz MC. Assemblytics: a web analytics tool for the detection of variants from an assembly. Bioinformatics. 2016 Oct 1;32(19):3021–3.

35. Boyer RS, Moore JS. A fast string searching algorithm. Commun ACM. 1977 Oct 1;20(10):762–72.

36. Barckmann B, El-Barouk M, Pélisson A, Mugat B, Li B, Franckhauser C, et al. The somatic piRNA pathway controls germline transposition over generations. Nucleic Acids Res. 2018 Oct 12;46(18):9524–36.

37. Kolmogorov M, Yuan J, Lin Y, Pevzner PA. Assembly of long, error-prone reads using repeat graphs. Nat Biotechnol. 2019 May;37(5):540–6.

38. Shafin K, Pesout T, Lorig-Roach R, Haukness M, Olsen HE, Bosworth C, et al. Nanopore sequencing and the Shasta toolkit enable efficient de novo assembly of eleven human genomes. Nat Biotechnol. 2020 Sep;38(9):1044–53.

39. Ruan J, Li H. Fast and accurate long-read assembly with wtdbg2. Nat Methods. 2020 Feb;17(2): 155–8.

40. Vaser R, Sović I, Nagarajan N, Šikić M. Fast and accurate de novo genome assembly from long uncorrected reads. Genome Res. 2017 Jan 5;27(5):737–46.

41. Alonge M, Soyk S, Ramakrishnan S, Wang X, Goodwin S, Sedlazeck FJ, et al. RaGOO: fast and accurate reference-guided scaffolding of draft genomes. Genome Biology. 2019 Oct 28;20(1):224.

42. Wick R. rrwick/Filtlong [Internet]. 2022 [cited 2022 Jul 20]. Available from: https://github.com/rrwick/Filtlong

43. Valiente-Mullor C, Beamud B, Ansari I, Francés-Cuesta C, García-González N, Mejía L, et al. One is not enough: On the effects of reference genome for the mapping and subsequent analyses of short-reads. PLoS Comput Biol. 2021 Jan 27;17(1):e1008678.

44. Solares EA, Chakraborty M, Miller DE, Kalsow S, Hall K, Perera AG, et al. Rapid Low-Cost Assembly of the Drosophila melanogaster Reference Genome Using Low-Coverage, Long-Read Sequencing. G3 (Bethesda). 2018 03;8(10):3143–54.

45. Gotea V, Veeramachaneni V, Makałowski W. Mastering seeds for genomic size nucleotide BLAST searches. Nucleic Acids Research. 2003 Dec 1;31(23):6935–41.

46. Camacho C, Coulouris G, Avagyan V, Ma N, Papadopoulos J, Bealer K, et al. BLAST+: architecture and applications. BMC Bioinformatics. 2009 Dec 15;10(1):421.

47. Sedlazeck FJ, Rescheneder P, Smolka M, Fang H, Nattestad M, Haeseler A von, et al. Accurate detection of complex structural variations using single-molecule sequencing. Nat Methods. 2018 Jun;15(6):461–8.

48. Li H, Handsaker B, Wysoker A, Fennell T, Ruan J, Homer N, et al. The Sequence Alignment/Map format and SAMtools. Bioinformatics. 2009 Aug 15;25(16):2078–9.

49. Li Y, Wang S, Bi C, Qiu Z, Li M, Gao X. DeepSimulator1.5: a more powerful, quicker and lighter simulator for Nanopore sequencing. Bioinformatics. 2020 Apr 15;36(8):2578–80.

50. Orjuela J, Comte A, Ravel S, Charriat F, Vi T, Sabot F, et al. CulebrONT: a streamlined long reads multi-assembler pipeline for prokaryotic and eukaryotic genomes [Internet]. bioRxiv; 2021 [cited 2022 Jul 20]. p. 2021.07.19.452922. Available from: https://www.biorxiv.org/content/10.1101/2021.07.19.452922v1

