## Supplemental Figure 1 for "TrEMOLO: Accurate transposable element allele frequency estimation using long-read sequencing data combining assembly and mapping-based approaches"

**Figure S1**

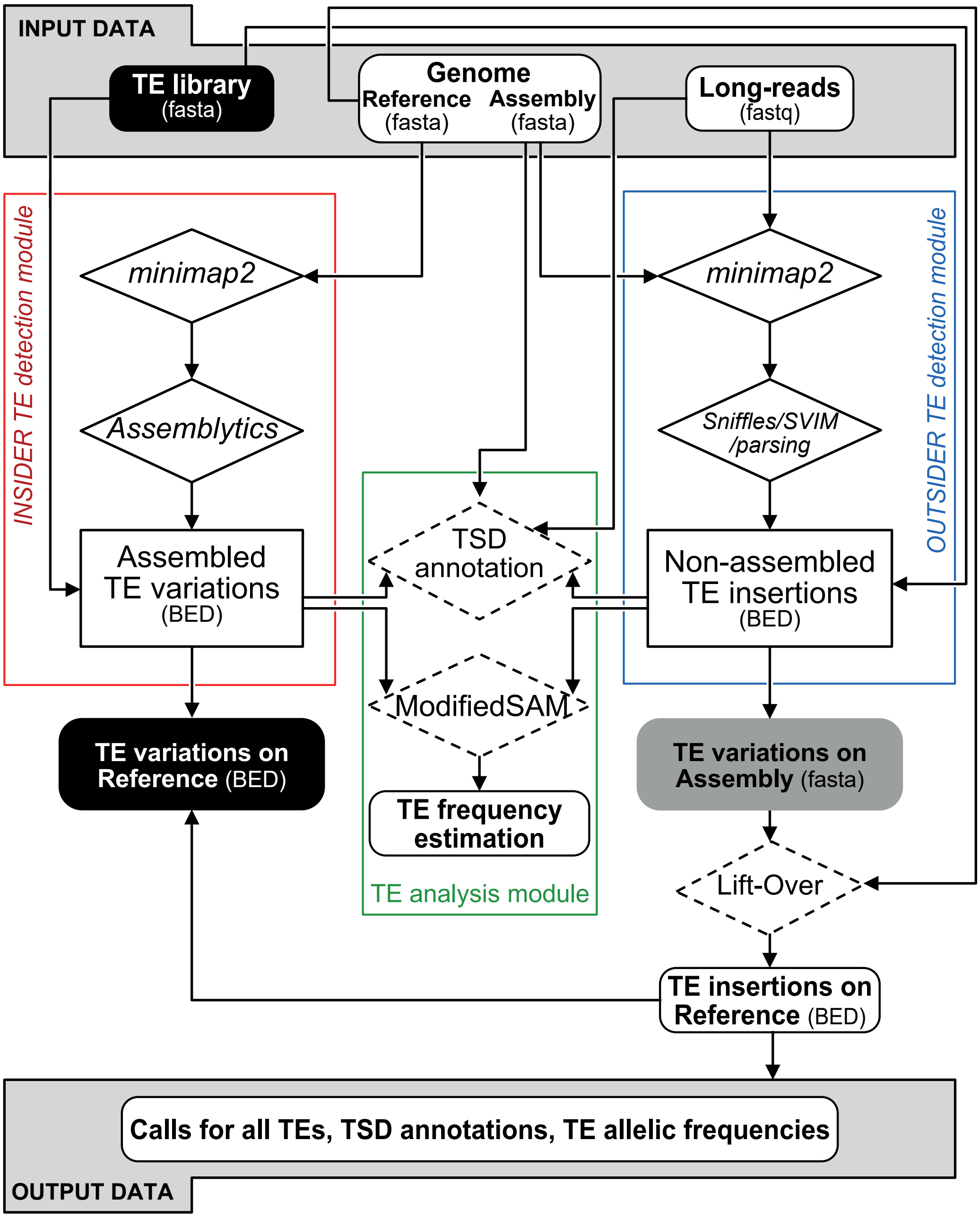


**Schematic representation of TrEMOLO method for TE detection**

The INSIDER TE module (red) detects TE insertions/deletions in genome assemblies and the OUTSIDER TE module detects TE insertions/deletions by mapping reads on these assemblies (blue). The TE analysis module allows the characterization of the TE insertions by determining the TSDs and estimating the TE frequencies. The coordinates of the insertions can be determined on the reference genome by lift-over. The final output files are calls for all TEs, TSD annotations and TE allelic frequencies.
