## Supplemental Figure 2 for "TrEMOLO: Accurate transposable element allele frequency estimation using long-read sequencing data combining assembly and mapping-based approaches"

Figure S2

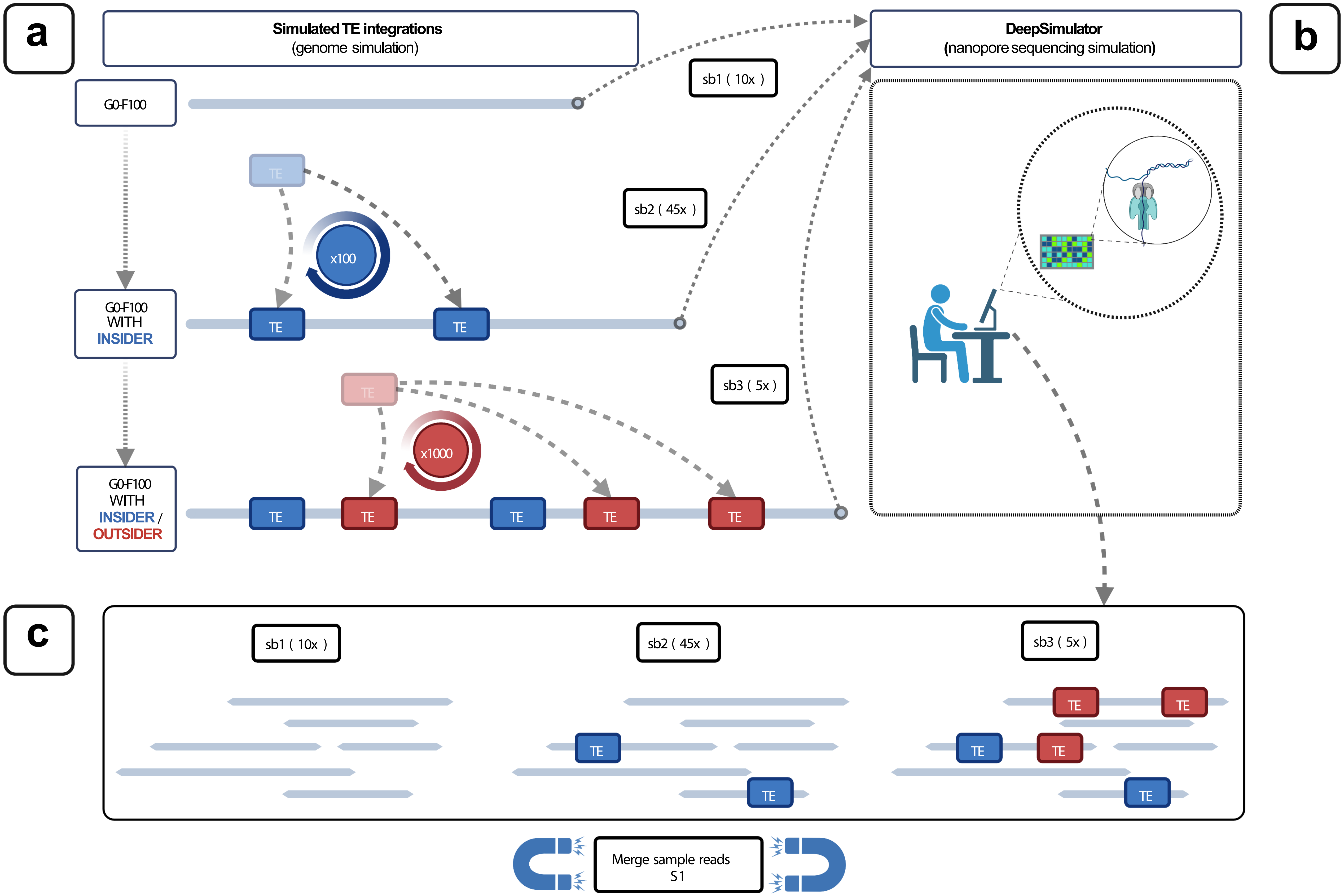


**Schematic representation to get benchmarking datasets**

a) Genome simulation using the G0-F100 unmasked assembled genome. b) read simulation using DeepSimulator. c) Proportion of the different subsamples in the S1 simulated reads.
