## Supplemental Figure 3 for "TrEMOLO: Accurate transposable element allele frequency estimation using long-read sequencing data combining assembly and mapping-based approaches"

**Figure S3**

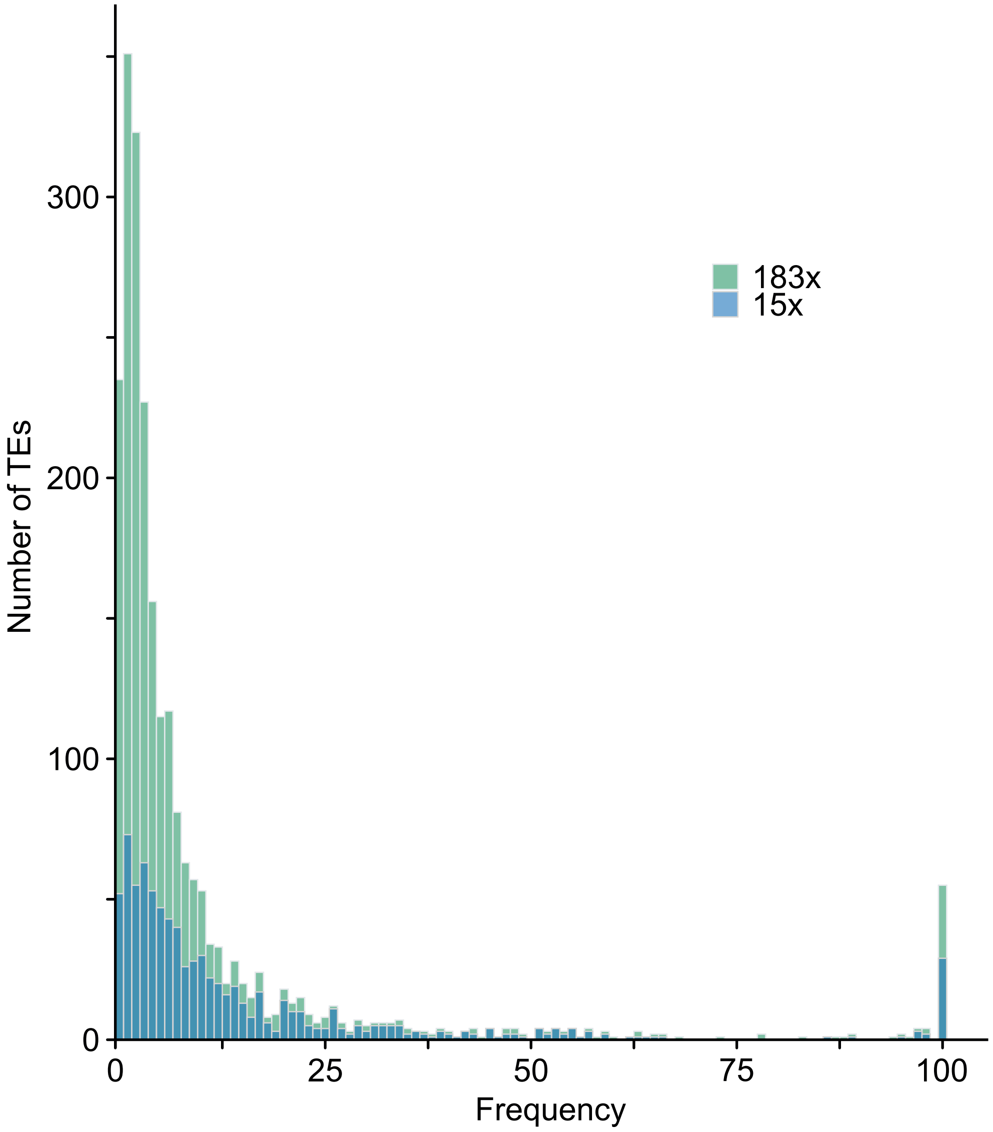


**Impact of down-sampling on low frequency TE detection**

Frequency distributions of OUTSIDER TE insertions detected in the G73 (183x) dataset (green) and in the 15x dataset (blue).
